# Architecture and self-assembly of the SARS-CoV-2 nucleocapsid protein

**DOI:** 10.1101/2020.05.17.100685

**Authors:** Qiaozhen Ye, Alan M.V. West, Steve Silletti, Kevin D. Corbett

**Author notes:** Correspondence should be addressed to Kevin D. Corbett: 9500 Gilman Drive, #3206, La Jolla, CA 92093.

## Abstract

The COVID-2019 pandemic is the most severe acute public health threat of the twenty-first century. To properly address this crisis with both robust testing and novel treatments, we require a deep understanding of the life cycle of the causative agent, the SARS-CoV-2 coronavirus. Here, we examine the architecture and self-assembly properties of the SARS-CoV-2 nucleocapsid protein, which packages viral RNA into new virions. We determined a 1.4 Å resolution crystal structure of this protein’s N2b domain, revealing a compact, intertwined dimer similar to that of related coronaviruses including SARS-CoV. While the N2b domain forms a dimer in solution, addition of the C-terminal spacer B/N3 domain mediates formation of a homotetramer. Using hydrogen-deuterium exchange mass spectrometry, we find evidence that at least part of this putatively disordered domain is structured, potentially forming an α-helix that self-associates and cooperates with the N2b domain to mediate tetramer formation. Finally, we map the locations of amino acid substitutions in the N protein from over 38,000 SARS-CoV-2 genome sequences. We find that these substitutions are strongly clustered in the protein’s N2a linker domain, and that substitutions within the N1b and N2b domains cluster away from their functional RNA binding and dimerization interfaces. Overall, this work reveals the architecture and self-assembly properties of a key protein in the SARS-CoV-2 life cycle, with implications for both drug design and antibody-based testing.

## Introduction

SARS-CoV-2 ^1,2^ is the third coronavirus to cross from an animal reservoir to infect humans in the 21^st^ century, after SARS (severe acute respiratory syndrome coronavirus) ^3,4^ and MERS (Middle-East respiratory syndrome coronavirus) ^5^. Isolation and sequencing of SARS-CoV-2 was reported in January 2020, and the virus was found to be highly related to SARS and share a probable origin in bats ^2,6^. Since its emergence in December 2019 in Wuhan, China, the virus has infected about 7 million people and caused nearly 400,000 deaths as of early June, 2020 (https://coronavirus.jhu.edu). The high infectivity of SARS-CoV-2 and the worldwide spread of this ongoing outbreak highlight the urgent need for public health measures and therapeutics to limit new infections. Moreover, the severity of the atypical pneumonia caused by SARS-CoV-2 (COVID-2019), often requiring multi-week hospital stays and the use of invasive ventilators ^7–9^, highlights the need for therapeutics to lessen the severity of individual infections.

Current therapeutic strategies against SARS-CoV-2 target major points in the life-cycle of the virus. The antiviral Remdesivir, first developed against Ebola virus ^10,11^, inhibits the viral RNA-dependent RNA polymerases of a range of coronaviruses including SARS-CoV-2 ^12–14^ and has shown promise against SARS-CoV-2 in small-scale trials in both primates and humans ^15,16^. Another target is the viral protease (Mpro/3CLpro), which is required to process viral polyproteins into their active forms ^17^. Finally, the transmembrane spike (S) glycoprotein mediates binding to host cells through the angiotensin converting enzyme 2 (ACE2) and transmembrane protease, serine 2 (TMPRSS2) proteins, and mediates fusion of the viral and host cell membranes ^18–21^. As the most prominent surface component of the virus, the spike protein is the major target of antibodies in patients, and is the focus of several current efforts at SARS-CoV-2 vaccine development. Initial trials using antibody-containing plasma of convalescent COVID-19 patients has also shown promise in lessening the severity of the disease ^22^.

While the above efforts target viral entry, RNA synthesis, and protein processing, there has so far been less emphasis on other steps in the viral life cycle. One critical step in coronavirus replication is the assembly of the viral genomic RNA and nucleocapsid (N) protein into a ribonucleoprotein (RNP) complex, which in betacoronaviruses like SARS-CoV-2 is thought to form a helical filament structure that is packaged into virions through interactions with the membrane-spanning membrane (M) protein ^23–28^. Despite its location within the viral particle rather than on its surface, patients infected with SARS-CoV-2 show higher and earlier antibody responses to the nucleocapsid protein than the surface spike protein ^29,30^. As such, a better understanding of the SARS-CoV-2 N protein’s structure, and structural differences between it and N proteins of related coronaviruses including SARS-CoV, may aid the development of sensitive and specific immunological tests.

Coronavirus N proteins possess a shared domain structure with an N-terminal RNA-binding domain and a C-terminal domain responsible for dimerization. The assembly of N protein dimers into higher-order complexes is not well understood, but likely involves cooperative interactions between the dimerization domain and other regions of the protein, plus the bound RNA ^31–39^. Here, we present a high-resolution structure of the SARS-CoV-2 N dimerization domain, revealing an intertwined dimer similar to that of related betacoronaviruses. We also analyze the self-assembly properties of the SARS-CoV-2 N protein, and show that higher-order assembly requires both the dimerization domain and the extended, disordered C-terminus of the protein. Together with other work revealing the structure and RNA-binding properties of the nucleocapsid N-terminal domain, these results lay the groundwork for a comprehensive understanding of SARS-CoV-2 nucleocapsid assembly and architecture.

## Results

### Structure of the SARS-CoV-2 Nucleocapsid dimerization domain

Betacoronavirus nucleocapsid (N) proteins share a common overall domain structure, with ordered RNA-binding (N1b) and dimerization (N2b) domains separated by short regions with high predicted disorder (N1a, N2a, and spacer B/N3; **Figure 1A**). Self-association of the full-length SARS-CoV N protein and the isolated C-terminal region (domains N2b plus spacer B/N3; residues 210-422) was first demonstrated by yeast two-hybrid analysis ^31^, and the purified full-length protein was shown to self-associate into predominantly dimers in solution ^32^. The structures of the N2b domain of SARS-CoV and several related coronaviruses confirmed the obligate homodimeric structure of this domain ^33–39^, and other work showed that the region C-terminal to this domain mediates further self-association into tetramer, hexamer, and potentially higher oligomeric forms ^40–42^. Other studies have suggested that the protein’s N-terminal region, including the RNA-binding N1b domain, can also self-associate ^43,44^, highlighting the possibility that assembly of full-length N into helical filaments is mediated by cooperative interactions among several interfaces.

**Figure 1.**
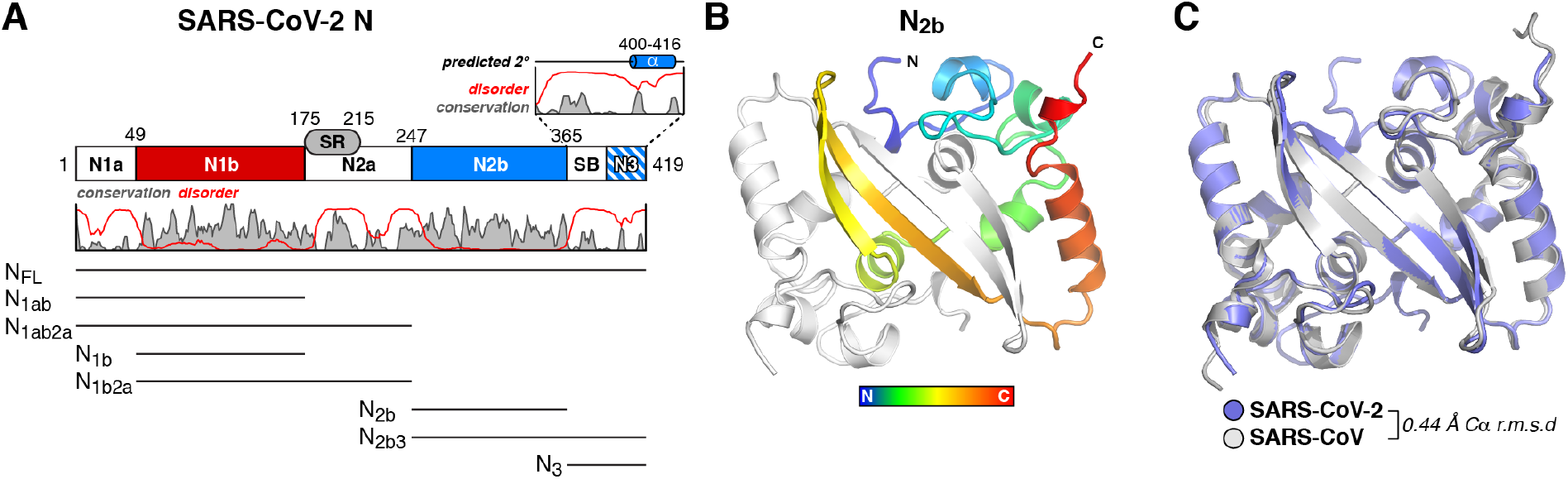
Structure of the SARS-CoV-2 Nucleocapsid dimerization domain. (A) Domain structure of the SARS-CoV-2 Nucleocapsid protein, as defined previously ^45,46^, with plot showing the Jalview alignment conservation score (three-point smoothed; gray) ^59^ and DISOPRED3 disorder propensity (red) ^60^ for nine related coronavirus N proteins (SARS-CoV, SARS-CoV-2, MERS-CoV, HCoV-OC43, HCoV-HKU1, HCoV-NL63, and HCoV-229E, IBV (Infectious Bronchitis virus), and MHV (Murine Hepatitis virus)). SR: serine/arginine rich domain; SB; spacer B. The boundary between SB and N3 is not well-defined due to low identity between SARS-CoV/SARS-CoV-2 and MHV N proteins ^46^. All purified truncations are noted at bottom. (B) Two views of the SARS-CoV-2 N_2b_ dimer, with one monomer colored as a rainbow (N-terminus blue, C-terminus red) and the other colored white. See **Figure S1A** for comparison with other structures of this domain. (C) Structural overlay of the SARS-CoV-2 N_2b_ dimer (blue) and the equivalent domain of SARS-CoV-N (PDB ID 2GIB) ^34^.

To characterize the structure and self-assembly properties of the SARS-CoV-2 nucleocapsid, we first cloned and purified the protein’s N2b dimerization domain (N_2b_; residues 247-364) ^45,46^. We crystallized and determined two high-resolution crystal structures of N_2b_; a 1.45 Å resolution structure of His_6_-tagged N_2b_ at pH 8.5, and a 1.42 Å resolution structure of N_2b_ after His_6_-tag cleavage, at pH 4.5 (see **Materials and Methods** and **Table S1**). These structures reveal a compact, tightly intertwined dimer with a central four-stranded β-sheet comprising the bulk of the dimer interface (**Figure 1B**). This interface is composed of two β-strands and a short α-helix from each protomer that extend toward the opposite protomer and pack against its hydrophobic core. The asymmetric units of both structures contain two N_2b_ dimers, giving four crystallographically independent views of the N_2b_ dimer. These four dimers differ only slightly, showing overall Cα r.m.s.d values of 0.15-0.19 Å and with most variation arising from loop regions (**Figure S1A**). Our structures also overlay closely with three other recently-deposited structures of the SARS-CoV-2 N2b domain (PDB IDs 6WJI, 6YUN, and 7C22; all unpublished). One of these structures (PDB ID 7C22) was crystallized in equivalent conditions as our structure of untagged N_2b_. Including all of these structures, there are now eight independent crystallographic views of the SARS-CoV-2 N2b domain dimer (16 total protomers) in four crystal forms at pH 4.5, 7.5, 7.8, and 8.5. All of these structures overlay closely, with an overall Cα r.m.s.d of 0.15-0.31 Å (**Figure S1A**).

The structure of N_2b_ closely resembles that of related coronaviruses, including SARS-CoV, Infectious Bronchitis Virus (IBV), MERS-CoV, and HCoV-NL63 ^33–38^. The structure is particularly similar to that of SARS-CoV, with which the N2b domain shares 96% sequence identity; only five residues differ between these proteins’ N2b domains (SARS-CoV Gln268 → SARS-CoV-2 A267, D291→E290, H335→Thr334, Gln346→Asn345, and Asn350→Gln349), and the structures are correspondingly similar with an overall Cα r.m.s.d of 0.44 Å across the N_2b_ dimer (**Figure 1C**).

A crystal structure of the SARS-CoV N protein revealed a helical assembly of N2b domain dimers that was proposed as a possible structure for the observed helical nucleocapsid filaments in virions ^33^. We therefore examined the packing of N2b domain dimers in the five crystal structures of SARS-CoV-2 N, four of which show distinct space groups and unit cell parameters. We identified two dimer-dimer packing modes that are observed in four or all five structures, respectively (**Figure S1B**). These packing modes do not result in the assembly of a helical filament, and do not strongly correlate with highly conserved surfaces on the N2b domain. This evidence, combined with our finding that N_2b_ forms solely dimers in solution (see below), suggests that packing of N2b domain dimers does not underlie higher-order assembly of SARS-CoV-2 N protein filaments.

### N protein variation in SARS-CoV-2 patient samples

Since the first genome sequence of SARS-CoV-2 was reported in January 2020 ^2,6^, over 38,000 full genomic sequences have been deposited in public databases (as of June 4, 2020). To examine the variability of the N protein in these sequences, we downloaded a comprehensive list of reported mutations within the SARS-CoV-2 N gene in a set of 38,318 genome sequences from the China National Center for Bioinformation, 2019 Novel Coronavirus Resource. Among these sequences, there are 10,983 instances of amino acid substitutions spread across 250 of the 419 amino acids of the N protein (**Figure 2A**, **Table S2**). These variants are strongly clustered in the N2a linker domain, particularly in the serine/arginine-rich subdomain (SR in **Figure 2A**). The most common substitutions are R203K and G204R, which occur together as the result of a common trinucleotide substitution in genomic positions 28881-28883, from GGG to AAC (~4,100 of the 38,318 sequences in our dataset; **Figure S2A-B**). While positions 203 and 204 accounted for over two-thirds of the total individual amino acid substitutions in this dataset, the N2a region shows a strong enrichment of mutations even when these positions are not considered (**Figure 2A**). In contrast to the enrichment of missense mutations in the N2a domain, synonymous mutations were distributed relatively equally throughout the protein (**Figure S2C**). Thus, these data suggest that the N2a domain is uniquely tolerant of mutations, in keeping with its likely structural role as a disordered linker between the RNA-binding N1b domain and the N2b dimerization domain.

**Figure 2.**
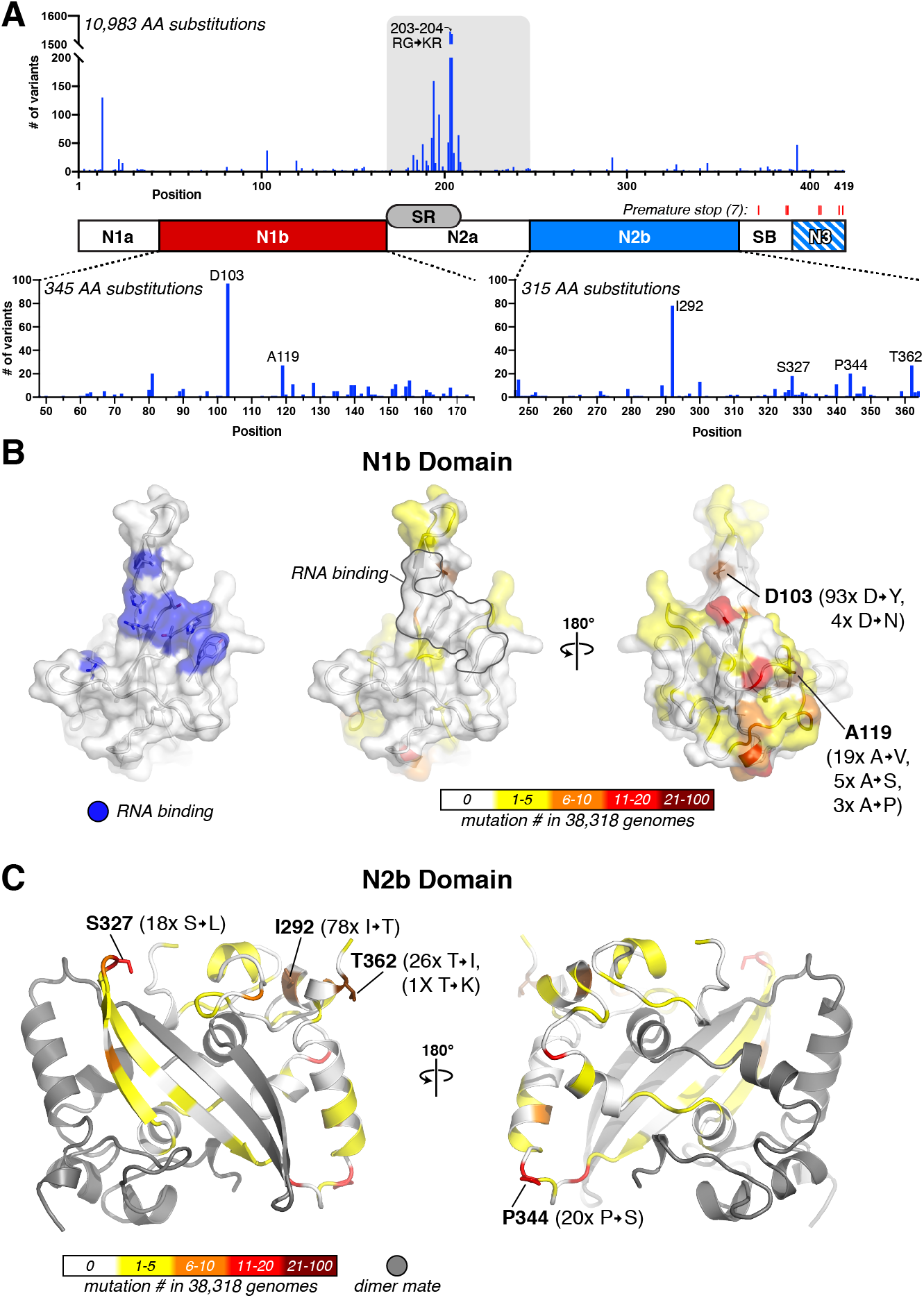
N protein variability in SARS-CoV-2 patient sequences. (A) *Top:* Plot showing the number of observed amino acid variants at each position in the N gene in 16,975 SARS-CoV-2 genomes (details in **Table S2**). The most highly-mutated positions are R203 and G204, which are each mutated more than 4,000 times due to a prevalent trinucleotide substitution (**Figure S2A-B**). Red tick marks indicate the locations of seven premature stop mutations detected (two sequences contained stop codons at residue 256; not graphed). *Bottom:* Plots showing amino acid variants in the N1b and N2b domains. (B) Surface views of the N protein N1b domain (PDB ID 6VYO; Center for Structural Genomics of Infectious Diseases (CSGID), unpublished). At left, blue indicates RNA-binding residues identified by NMR peak shifts (A50, T57, H59, R92, I94, S105, R107, R149, and Y172) ^48^. At right, two views colored by the number of variants at each position observed in a set of 38,318 SARS-CoV-2 genomes. The two most frequently-mutated residues are shown in stick view and labeled. Only one mutation (A50E, observed in one sequence) overlaps the putative RNA binding surface. (C) Cartoon view of the N protein N2b domain, with one monomer colored gray and the other colored by the number of variants at each position observed in a set of 38,318 SARS-CoV-2 genomes. The four most frequently-mutated residues are shown in stick view and labeled.

While the majority of N protein mutations are in the N2a domain, we nonetheless identified 345 instances of amino acid variants in the RNA-binding N1b domain, and 315 instances in the N2b domain. We mapped these onto high-resolution structures of both domains (**Figure 2B-C**). Two high-resolution crystal structures of the SARS-CoV-2 N1b domain have been determined (PDB ID 6M3M and 6VYO) ^47^, and a recent NMR study determined a solution structure of the domain and defined its likely RNA binding surface (**Figure 2B**) ^48^. In keeping with its functional importance, the identified RNA binding surface shows only a single mutation in this dataset (**Figure 2B**; middle panel). In the N2b domain, most mutations occur on surface residues, particularly in loop regions, while the functionally-important dimer interface is largely invariant (**Figure 2C**).

Finally, the 38,318 SARS-CoV-2 genome sequences contain nine sequences with reported nonsense/premature stop codons in the N protein. Two of these are located at position 256 within the N2b domain, while the remaining seven are located in the spacer B/N3 region between positions 372-418 (**Figure 2A**).

### Self-association of the SARS-CoV-2 N protein

Our structures of the SARS-CoV-2 N protein N2b domain reveal that, as in related coronaviruses, this domain mediates homodimer formation. We next systematically investigated the molecular basis for higher-order self-assembly of the SARS-CoV-2 nucleocapsid. We first purified the full-length N protein (N_FL_) for analysis of its oligomeric state. While our initial attempts at purification yielded large aggregates significantly contaminated with nucleic acid (**Figure S3A**), purification of the protein in high-salt buffer (1M NaCl) and in the presence of both DNase and RNase yielded pure N_FL_ (**Figure S3B**). Using size exclusion chromatography coupled to multi-angle light scattering (SEC-MALS), we determined that NFL on its own assembles into a homotetramer in solution (**Figure 3A**).

**Figure 3.**
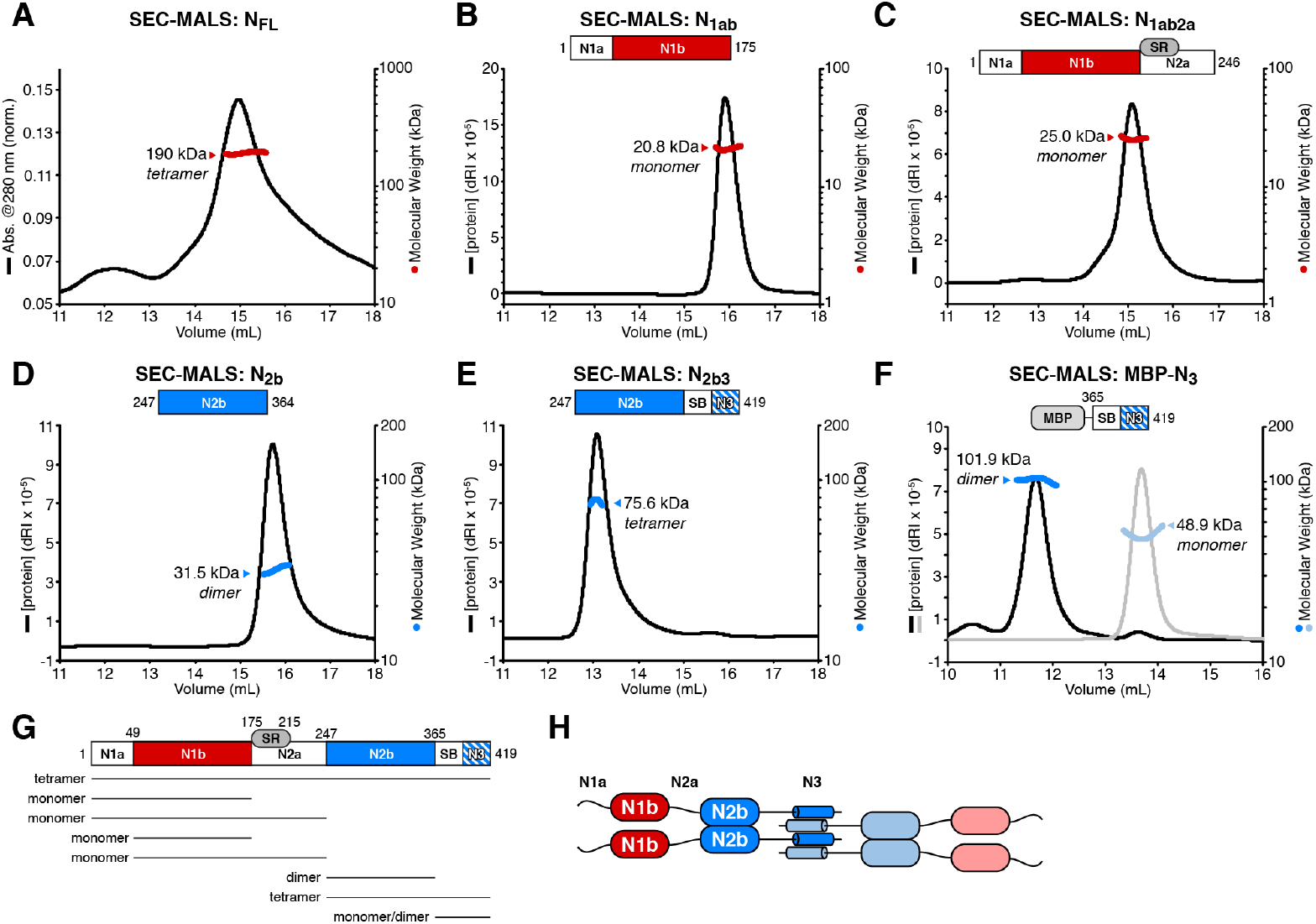
The C-terminus of N mediates tetramer formation. (A) Size exclusion chromatography (Superose 6) coupled to multi-angle light scattering (SEC-MALS) analysis of full-length SARS-CoV-2 N. The measured MW of 190.0 kDa closely matches that of a tetramer (182.5 kDa). See **Figure S3B** for SDS-PAGE analysis of all purified proteins. (B) Size exclusion chromatography (Superdex 200 Increase; used for panels B-F) coupled to multi-angle light scattering (SEC-MALS) analysis of SARS-CoV-2 N_1ab_ (residues 2-175). The measured MW of 20.8 kDa closely matches that of a monomer (18.9 kDa). dRI: differential refractive index. (C) SEC-MALS analysis of SARS-CoV-2 N_1ab2a_ (residues 2-246). The measured MW of 25.0 kDa is slightly less than that of a monomer (26.2 kDa), reflecting partial proteolysis within the N2a domain (**Figure S3B**). (D) SEC-MALS analysis of SARS-CoV-2 N_2b_. The measured MW (31.5 kDa) closely matches that of a homodimer (26.5 kDa). (E) SEC-MALS analysis of SARS-CoV-2 N_2b3_. The measured MW (75.6 kDa) closely matches that of a homotetramer (77.4 kDa). (F) SEC-MALS analysis of MBP-SARS-CoV-2 N_3_ (“peak 1” black/dark blue; “peak 2” gray/light blue) The measured MW of peak 1 (101.9 kDa) and peak 2 (48.9 kDa) closely match those of a homodimer (101.7 kDa) and a monomer (50.9 kDa). The small peak at 10.5 mL suggests higher-order self-assembly. (G) Schematic summary of size exclusion and SEC-MALS results on N protein constructs. See **Figure S3C-D** for SEC-MALS analysis of N_1b_ (residues 49-174) and N_1b2a_ (residues 49-246). (H) Proposed self-assembly mechanism of SARS-CoV-2 N. Dimerization is mediated by the N2b domain, and these dimers self-associate through the N3 region to form homotetramers.

To determine the molecular basis for homotetramer assembly, we purified a series of truncation constructs encompassing the ordered N1b and N2b domains and their associated linker domains (N1a, N2a, and spacer B/N3; **Figure 1A**). We characterized four truncations encompassing the protein’s N-terminal regions, including N_1ab_ (residues 2-175), N_1b_ (residues 49-175), N_1ab2a_ (residues 2-246), and N_1b2a_ (residues 49-246). All four of these truncations are monomeric in solution as determined by SEC-MALS (**Figure 3B-C, Figure S3C-D**). We next analyzed N2b, which forms a homodimer in our crystal structures. As expected, N2b is dimeric in solution (**Figure 3D**).

Finally, we analyzed the contribution of the C-terminal spacer B/N3 region to N protein self-assembly. Prior work with the Murine Hepatitis Virus (MHV) N protein showed that this region can, on its own, incorporate into nucleocapsid structures that lack the associated Membrane (M) protein, suggesting that the region mediates a homotypic interaction between N proteins ^46^. Other work with SARS-CoV and HCoV-229E N proteins also found that the C-terminal spacer B/N3 region is required for higher-order assembly of tetramers and larger oligomers ^40–42^. We purified a construct encoding N2b and the spacer B/N3 region (N_2b3_, residues 247-419) and found that it forms a homotetramer (**Figure 3E**). We also analyzed self-assembly of the spacer B/N3 region on its own by performing SEC-MALS analysis on this isolated region (N_3_, residues 365-419) fused to a His_6_-maltose binding protein (MBP) tag. Initial purification of His_6_-MBP-N_3_ yielded two peaks on the final size exclusion column, which we separated pooled and analyzed by SEC-MALS. We found that these two peaks correspond to a monomer and a dimer, respectively (**Figure 3F**). The pooled dimer population also showed a small population of potentially higher-order oligomers (**Figure 3F**). Together, these data suggest that assembly of betacoronavirus N protein filaments likely proceeds through at least three steps, each mediated by different oligomerization interfaces: (1) dimerization mediated by the N2b domain; (2) tetramerization mediated by the spacer B/N3 region (**Figure 3G-H**); and (3) oligomer/filament assembly mediated by cooperative RNA binding and potential higher-order self-association of N homotetramers.

To gain structural insight into how the spacer B/N3 region mediates N protein self-association, we performed hydrogen-deuterium exchange mass spectrometry (HDX-MS) on N_2b_ and N_2b3_ (**Figure 4**). By probing the rate of exchange of amide hydrogen atoms with deuterium atoms in a D_2_O solvent, HDX-MS provides information on the level of secondary structure and solvent accessibility across an entire protein. We found that H-D exchange rates within N_2b_ largely agreed with our crystal structure: regions in β-strands or α-helices showed low exchange rates consistent with high order, while loop regions showed increased exchange rates consistent with their likely flexibility (**Figure 4A, C, D**).

**Figure 4.**
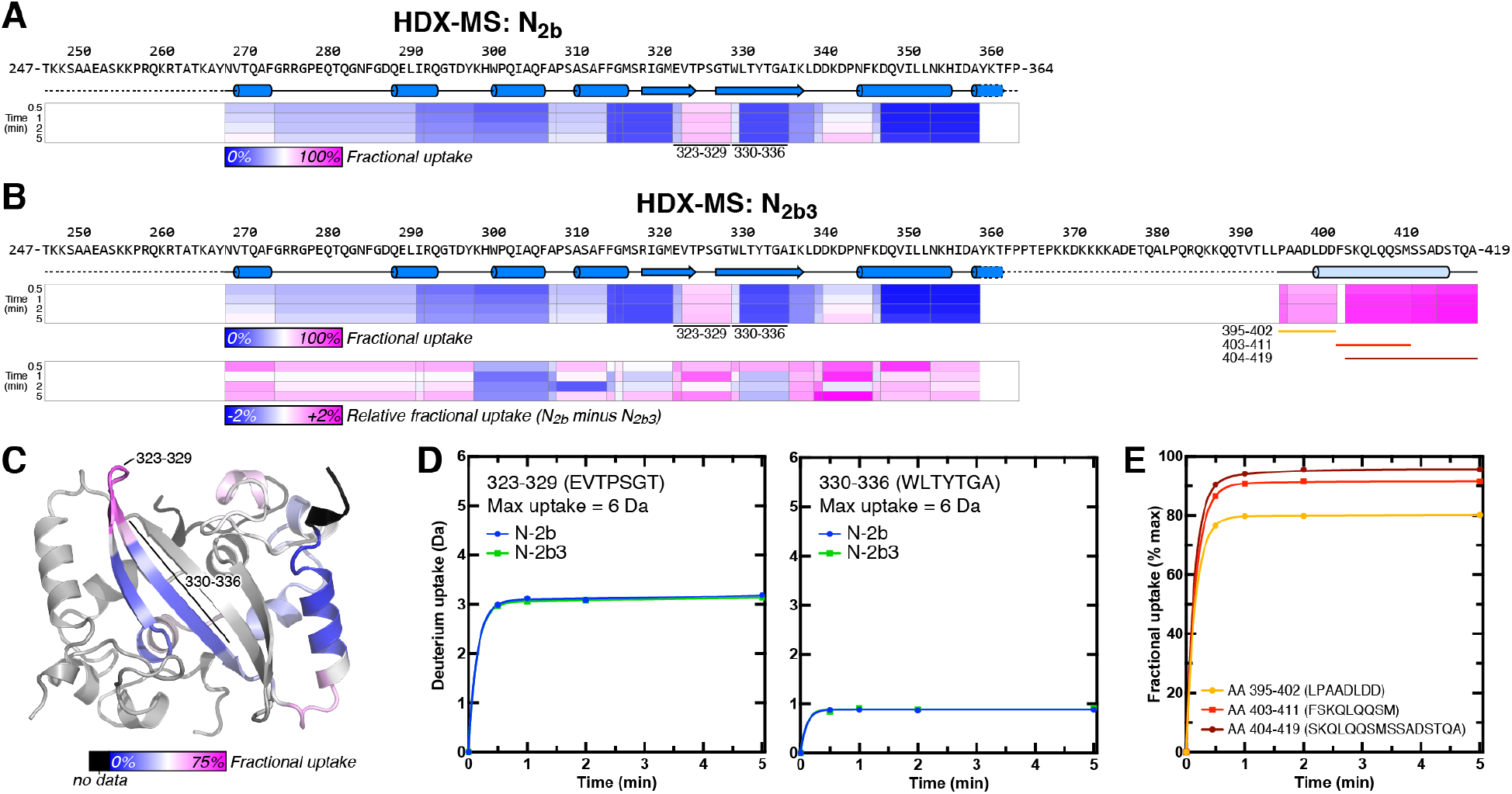
HDX-MS analysis of N_2b_ and N_2b3_. (A) Schematic showing the N_2b_ sequence and structure, plus protein regions detected by HDX-MS. Each peptide is colored by its fractional deuterium uptake during the course of the experiment (blue-white-magenta = 0-100% fractional uptake). (B) Schematic showing the N_2b3_ sequence and inferred structure (the α-helix spanning residues 400-416 is predicted by PSI-PRED), plus protein regions detected by HDX-MS. Two sets of exchange rates are shown: fractional deuterium uptake in N_2b3_ (upper box) colored as in panel A, and relative uptake comparing N_2b_ and N_2b3_ (lower box). (C) Structure of the N_2b_ dimer, with one monomer colored by fractional deuterium uptake (blue-white-magenta = 0-75% fractional uptake). (D) Uptake plots for two peptides within the ordered N2b domain, with uptake in N_2b_ indicated in blue and uptake in N_2b3_ indicated in green. The peptide covering residues 323-329 (located within a loop) is relatively exposed, while the peptide covering residues 330-336 (within a β-strand) is strongly protected from H-D exchange. (E) Uptake plots for three peptides in the C-terminal region of N_2b3_, plotted by fractional deuterium uptake. Peptides covering residues 395-402 (yellow) and 403-411 (red) show more protection than residues 404-419, suggesting that this region is partially structured. See **Figure S4A** for each peptide plotted by absolute deuterium uptake.

Compared to N_2b_, N_2b3_ contains an additional 56 amino acids (residues 365-419). While residues 360-394 were not detected in our HDX-MS analysis, we detected spectra for seven overlapping peptides spanning residues 395-419 at the protein’s extreme C-terminus (**Figure 4B**). While all of these peptides exhibited higher levels of exchange than the ordered N2b domain, peptides spanning the N-terminal part of this region (particularly residues 395-402) showed a degree of protection compared to those at the extreme C-terminus (residues 404-419; **Figure 4E**). This finding suggests that at least part of the spacer B/N3 domain possesses secondary structure and may mediate N_2b3_ tetramer formation. Indeed, analysis by the PSI-PRED server ^49^ suggests that this region may adopt an α-helical conformation (**Figure 1A**).

We next compared HDX-MS exchange rates of N_2b_ versus N_2b3_ for peptides within the N2b domain. We reasoned that if the C-terminus of N_2b3_ mediates tetramer formation, it may do so by docking against a surface in the N2b domain, which may be detectable by reduced deuterium uptake in the involved region. Contrary to this expectation, we found that the H-D exchange rates within the N2b domain were nearly identical between the two constructs, varying at most 2% in fractional deuterium uptake in individual peptides (**Figure 4B, D**). While these data do not rule out the possibility that the spacer B/N3 region docks against N2b, they nonetheless support our SEC-MALS data showing that spacer B/N3 independently self-associates to mediate N protein tetramer formation. Further supporting this idea, His_6_-MBP-N_3_ did not detectably bind N_2b_ in an in vitro pulldown assay using purified proteins (**Figure S4B**).

## Discussion

Given the severity of the ongoing COVID-19 pandemic, a deep understanding of the SARS-CoV-2 life cycle is urgently needed. Here, we examine the architecture and self-assembly properties of the SARS-CoV-2 nucleocapsid protein, a key player in viral replication responsible for packaging viral RNA into new virions. Through two high-resolution crystal structures, we show that this protein’s N2b domain forms a compact, strand-swapped dimer similar to that of related betacoronaviruses. While the N2b domain mediates dimer formation, we find that addition of the C-terminal spacer B/N3 domain mediates formation of a robust homotetramer. Finally, the full-length N protein assembles into tetramers on its own and large oligomers in the presence of nucleic acid, likely assembling through cooperative protein-protein interactions involving several regions of the protein, plus interactions mediated by nucleic acids.

Given the importance of nucleocapsid-mediated RNA packaging to the viral life cycle, small molecules that inhibit nucleocapsid self-assembly or mediate aberrant assembly may be effective at reducing the severity of infections and the infectivity of patients. The high resolution of our crystal structures will enable their use in virtual screening efforts to identify such nucleocapsid assembly modulators. Given the high conservation of the N2b domain in betacoronaviruses, these assembly modulators may also be effective at countering related viruses including SARS-CoV. As SARS-CoV-2 is unlikely to be the last betacoronavirus to jump from an animal reservoir to humans, the availability of such treatments could drastically alter the course of future epidemics.

The SARS-CoV-2 genome has been subject to unprecedented scrutiny, with over 38,000 individual genome sequences deposited in public databases as of early June, 2020. We used this set of genome sequences to identify over 10,000 instances of amino acid substitutions in the N protein, and showed that these variants are strongly clustered in the protein’s N2a linker domain. The ~650 substitutions we identified in the N1b and N2b domains were clustered away from these domains’ RNA binding and dimerization interfaces, reflecting the functional importance of these surfaces. Curiously, the identification of several nonsense mutations in the protein’s spacer B/N3 region suggests that this region may not be absolutely required for viral replication; this idea remains to be experimentally validated.

Given the early and strong antibody responses to the nucleocapsid displayed by SARS-CoV-2 infected patients, the distribution of mutations within this protein should be carefully considered as antibody-based tests are developed. The high variability of the N2a domain means that individual patient antibodies targeting this domain may not be reliably detected with tests using the reference N protein; especially if these antibodies recognize residues 203 and 204, which are mutated in a large fraction of infections. At the same time, patient antibodies targeting the conserved N1b and N2b domains may in fact cross-react with nucleocapsids of related coronaviruses like SARS-CoV. The availability of a panel of purified N protein constructs now makes it possible to systematically examine the epitopes of both patient-derived and commercial anti-nucleocapsid antibodies.

## Materials and Methods

### Cloning and Protein Purification

SARS-CoV-2 N protein constructs (N_FL_ (residues 2-419), N_1ab_ (2-175), N_1ab2a_ (2-246), N_1b_ (49-175), N_1b2a_ (49-246), N_2b_ (247-364), N_2b3_ (247-419)) were amplified by PCR from the IDT 2019-nCoV N positive control plasmid (IDT cat. # 10006625; NCBI RefSeq YP_009724397) and inserted by ligation-independent cloning into UC Berkeley Macrolab vector 2B-T (AmpR, N-terminal His_6_-fusion; Addgene #29666) for expression in *E. coli*. N_3_ (residues 365-419) was similarly inserted into UC Berkeley Macrolab vector 2C-T (AmpR, N-terminal His_6_-MBP fusion; Addgene #29706). Plasmids were transformed into *E. coli* strain Rosetta 2(DE3) pLysS (Novagen), and grown in the presence of ampicillin and chloramphenical to an OD_600_ of 0.8 at 37°C, induced with 0.25 mM IPTG, then grown for a further 16 hours at 18°C prior to harvesting by centrifugation. Harvested cells were resuspended in buffer A (25 mM Tris-HCl pH 7.5, 5 mM MgCl_2_ 10% glycerol, 5 mM β-mercaptoethanol, 1 mM NaN_3_) plus 500 mM NaCl and 5 mM imidazole pH 8.0. For purification, cells were lysed by sonication, then clarified lysates were loaded onto a Ni^2+^ affinity column (Ni-NTA Superflow; Qiagen), washed in buffer A plus 300 mM NaCl and 20 mM imidazole pH 8.0, and eluted in buffer A plus 300 mM NaCl and 400 mM imidazole. For cleavage of His_6_-tags, proteins were buffer exchanged in centrifugal concentrators (Amicon Ultra, EMD Millipore) to buffer A plus 300 mM NaCl and 20 mM imidazole, then incubated 16 hours at 4°C with TEV protease ^50^. Cleavage reactions were passed through a Ni^2+^ affinity column again to remove uncleaved protein, cleaved His_6_-tags, and His_6_-tagged TEV protease. Proteins were concentrated in centrifugal concentrators and purified by size-exclusion chromatography (Superdex 200; GE Life Sciences) in gel filtration buffer (25 mM Tris-HCl pH 7.5, 300 mM NaCl, 5 mM MgCl_2_, 10% glycerol, 1 mM DTT). Purified proteins were concentrated and stored at 4°C for analysis.

### SEC-MALS

For size exclusion chromatography coupled to multi-angle light scattering (SEC-MALS), 100 μL purified proteins at 2-5 mg/mL were injected onto a Superdex 200 Increase 10/300 GL column (GE Life Sciences) in gel filtration buffer. Light scattering and refractive index profiles were collected by miniDAWN TREOS and Optilab T-rEX detectors (Wyatt Technology), respectively, and molecular weight was calculated using ASTRA v. 6 software (Wyatt Technology).

### HDX-MS

H-D exchange experiments were conducted with a Waters Synapt G2S system. 5 μL samples containing 10 μM protein in gel filtration buffer were mixed with 55 μL of the same buffer made with D_2_O for several deuteration times (0 sec, 1 min, 2 min, 5 min, 10 min) at 15°C. The exchange was quenched for 2 min at 1°C with an equal volume of quench buffer (3M guanidine HCl, 0.1% formic acid). Proteins were cleaved with pepsin and separated by reverse-phase chromatography, then directed into a Waters SYNAPT G2s quadrupole time-of-flight (qTOF) mass spectrometer. Peptides were identified using PLGS version 2.5 (Waters, Inc.), deuterium uptake was calculated using DynamX version 2.0 (Waters Corp.), and uptake was corrected for back-exchange using DECA ^51^. Uptake plots were generated in Prism version 8.

### Crystallization and Structure Determination

For crystallization of untagged N_2b_, protein (40 mg/mL) in crystallization buffer (25 mM Tris-HCl pH 7.5, 200 mM NaCl, 5 mM MgCl_2_, and 1 mM Tris(2-carboxyethyl)phosphine) was mixed 1:1 with well solution containing 100 mM sodium acetate pH 4.5, 50 mM sodium/potassium tartrate, and 34% polyethylene glycol (PEG) 3350 at 20°C in hanging drop format. For crystallization of His_6_-tagged N_2b_, protein (40 mg/mL) in crystallization buffer was mixed 1:1 with well solution containing 100 mM Tris-HCl pH 8.5, 50 mM Ammonium Sulfate, and 38% PEG 3350 at 20°C in hanging drop format. Both untagged and His_6_-tagged N_2b_ formed large shard-like crystals, and were frozen in liquid nitrogen directly from the crystallization drop without further cryoprotection.

Diffraction data were collected at beamline 24ID-E at the Advanced Photon Source. Diffraction datasets were processed with the RAPD pipeline (https://github.com/RAPD/RAPD/), which uses XDS ^52^ for indexing and data reduction, and the CCP4 programs AIMLESS ^53^ and TRUNCATE ^54^ for scaling and conversion to structure factors. The structure of untagged N_2b_ was determined by molecular replacement in PHASER ^55^ using a dimer of the SARS-CoV N2b domain (PDB ID 2GIB) ^34^ as a template. The resulting model was manually rebuilt in COOT ^56^ and refined in phenix.refine ^57^ using positional, isotropic B-factor, and TLS (one group per chain) refinement. The structure of His_6_-tagged N2b was determined by molecular replacement from the structure of untagged N_2b_, then manually rebuilt and refined as above. Data collection statistics, refinement statistics, and database accession numbers for diffraction data and final structures can be found in **Table S1**. All structural figures were generated with PyMOL version 2.3.

### Nickel pulldown

Nickel pulldown assays were performed in buffer A plus 300 mM NaCl and 10 mM imidazole pH 8.0. Ten μg bait (His_6_-MBP-N_3_) was mixed with equal weights of each prey protein in 50 μl total reaction volume and incubated on ice for 30 minutes (5 μl “load” sample removed). 30 μl of a 50% slurry of Ni-NTA Superflow beads (Qiagen) was added and the mixture was incubated with occasional mixing on ice for 30 minutes. Beads were washed three times with 1 mL buffer, then bound proteins were eluted with the addition of 30 μl buffer A plus 300 mM NaCl and 250 mM imidazole pH 8.0. Ten μl of each eluate was analyzed by SDS-PAGE alongside the load samples.

### Bioinformatics

To examine variation in SARS-CoV-2 sequences, we downloaded a list of variants in the N gene in 38,318 genome sequences from China National Center for Bioinformation, 2019 Novel Coronavirus Resource (https://bigd.big.ac.cn/ncov?lang=en; downloaded June 3, 2020). We tabulated all mis-sense and nonsense mutations, and graphed the number of amino acid variants at each codon in Prism version 8 (all variant frequencies are listed in **Table S2**). To examine the prevalence of the trinucleotide substitution at genome positions 28881-28883, we downloaded a set of 200 SARS-CoV-2 genomes from the NCBI Virus Resource: https://www.ncbi.nlm.nih.gov/labs/virus/vssi/#/virus?SeqType_s=Nucleotide&VirusLineage_ss=SARS-CoV-2,%20taxid:2697049). We selected genomes with and without the substitution to show in **Figure S2A**. We used the NextStrain database ^58^ to visualize the prevalence of the N protein G204R mutation, which is diagnostic of the GGG→AAC trinucleotide substitution in positions 28881-28883. To visualize SARS-CoV-2 clade identity, we used the URL “https://nextstrain.org/ncov/global?c=clade_membership&l=unrooted”. To color by N protein residue 204 identity, we used the URL “https://nextstrain.org/ncov/global?c=gt-N_204&l=unrooted” (screenshots taken June 2, 2020).

## Acknowledgements

The authors thank the staff of Advanced Photon Source NE-CAT beamline 24ID-E for assistance with diffraction data collection, E. Komives for advice on HDX-MS interpretation, R. Lumpkin for assistance with the DECA software, and J. Pogliano, M. Daugherty, A. Schmidt, and members of the Corbett lab for helpful discussions. K.D.C. acknowledges generous institutional support from UC San Diego. The Waters Synapt HDX-MS at the UCSD BPMS Facility is supported by NIH S10 OD016234. The Northeastern Collaborative Access Team beamlines at the Advanced Photon Source are funded by the National Institute of General Medical Sciences from the National Institutes of Health (P30 GM124165). The Eiger 16M detector on the 24-ID-E beam line is funded by a NIH-ORIP HEI grant (S10OD021527). The Advanced Photon Source is a U.S. Department of Energy (DOE) Office of Science User Facility operated for the DOE Office of Science by Argonne National Laboratory under Contract No. DE-AC02-06CH11357.

## Supplemental Figures

**Figure S1.**
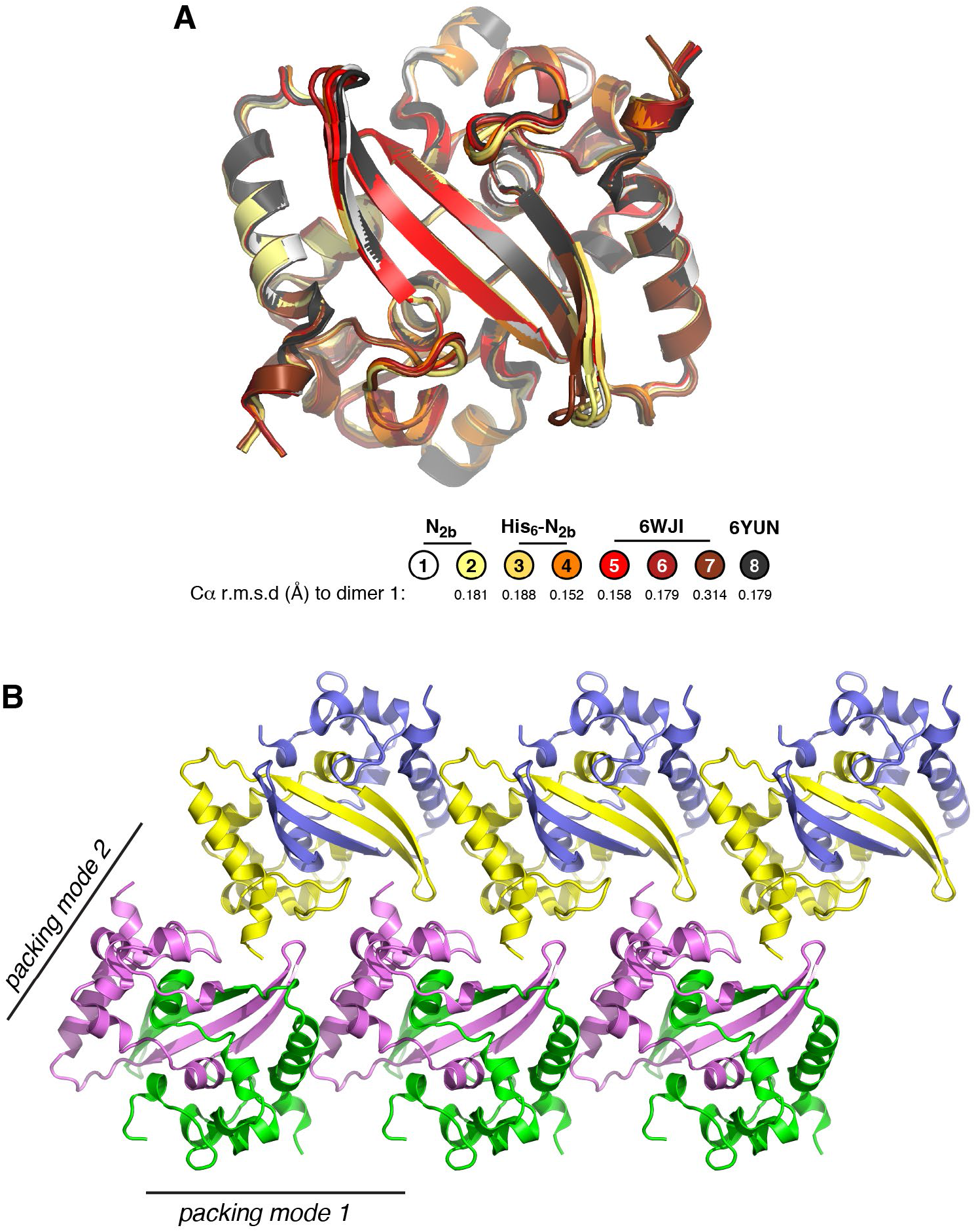
Comparison of SARS-CoV-2 N2b domain structures. (A) Overlay of eight independent views of the SARS-CoV-2 N2b domain dimer, including the two structures determined in this work (dimer 1=untagged N_2b_ chains A+B; dimer 2=untagged N_2b_ chains C+D, dimer 3=His_6_-tagged N_2b_ chains A+B, dimer 4=His_6_-tagged N_2b_ chains C+D), and two recently-deposited structures of the same domain (PDB ID 6WJI dimer 5=chains A+B, dimer 6=chains C+D, dimer 7=chains E+F; 6YUN dimer 8). A third deposited structure (PDB ID 7C22) has equivalent crystal packing as our structure of untagged N_2b_, so is not included in this analysis. The overall Cα r.m.s.d values of all dimers overlaid on dimer 1 are shown at bottom. (B) Crystal packing interactions in our structure of untagged N_2b_ (PDB ID 6WZO), showing two crystal packing modes for the SARS-CoV-2 N2b domain dimer. Of the five crystal structures of this domain (PDB IDs 6WJI, 6WZO, 6WZQ, 6YUN, and 7C22), all five show packing mode 1, and four (excluding 6WZQ) show packing mode 2.

**Figure S2.**
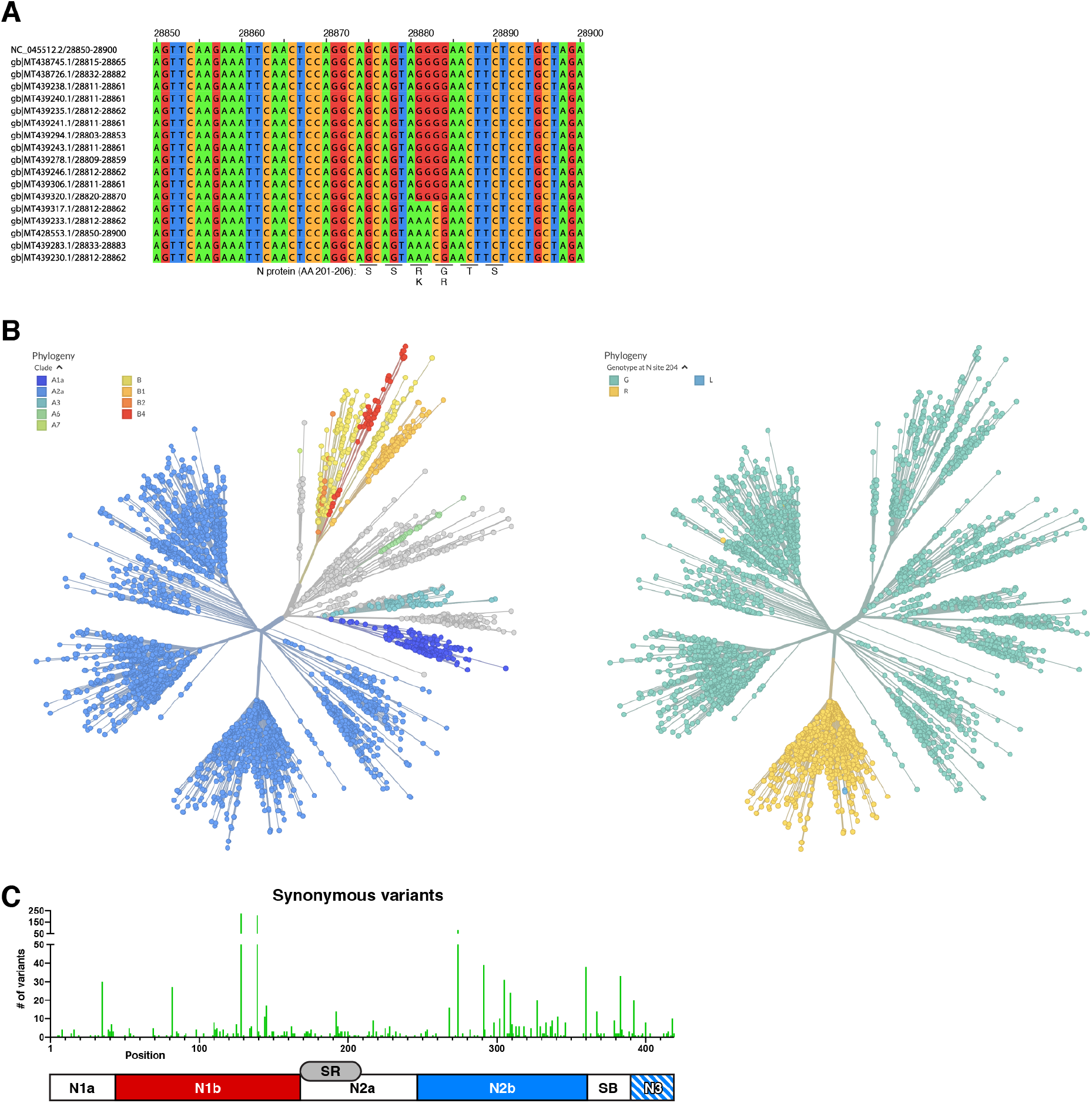
Mutations in sequenced SARS-CoV-2 isolates. (A) Sequence alignment of the SARS-CoV-2 reference genome (NCBI RefSeq NC_045512) and 17 selected sequences (from the NCBI Virus Resource; see **Materials and Methods**). Five genomes show the GGG→AAC trinucleotide substitution in position 28881-28883. Overall, roughly 10-15% of sequenced SARS-CoV-2 genomes show this trinucleotide substitution. (B) Screenshots from the Nextstrain resource ^58^ showing an unrooted tree of SARS-CoV-2 genomes colored by clade assignment (left) or by colored by N protein residue 204 identity (right). The G204R mutation (yellow) present in a large fraction of SARS-CoV-2 clade A2a samples is diagnostic of the GGG→AAC trinucleotide substitution in position 28881-28883. (C) Plot showing the number of observed synonymous variants at each position in the N gene in 38,318 SARS-CoV-2 genomes (details in **Table S2**).

**Figure S3.**
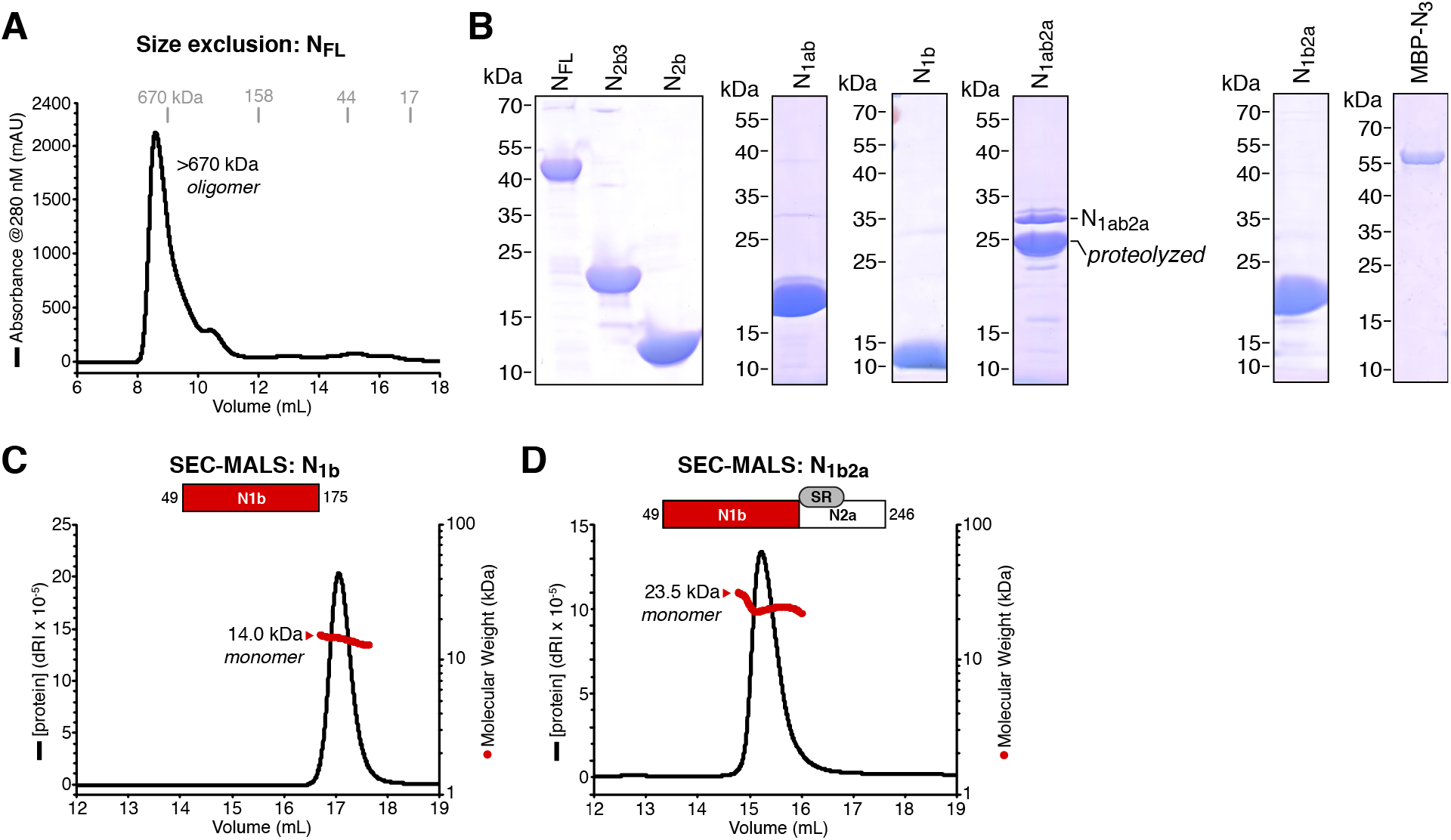
Purification and characterization of N protein constructs. (A) Size exclusion chromatography (Superdex 200 Increase) of N_FL_ protein purified with contaminating nucleic acid, as indicated by an A_260_/A_280_ ratio of 1.6. When bound to nucleic acid, N_FL_ assembles into a large oligomer. (B) SDS-PAGE analysis of purified SARS-CoV-2 N protein constructs: N_FL_ (residues 2-419), N_1ab_ (2-175), N_1ab2a_ (2-246), N_2b_ (247-364), N_2b3_ (247-419), N_1b_ (49-174), N_1b2a_ (49-246), and maltose binding protein (MBP)-fused N_3_ (364-419). N_1ab2a_ shows evidence of proteolytic cleavage within the N2a region during purification. N1b2a also possesses the N2a domain and shows evidence of proteolytic cleavage, but we could separate the cleaved and uncleaved fragments for analysis. (C) SEC-MALS analysis of SARS-CoV-2 N_1b_ (residues 49-175). The measured MW of 14.0 kDa closely matches that of a monomer (13.9 kDa). (D) SEC-MALS analysis of SARS-CoV-2 N_1b2a_ (residues 49-246). The measured MW of 23.5 kDa closely matches that of a monomer (21.1 kDa).

**Figure S4.**
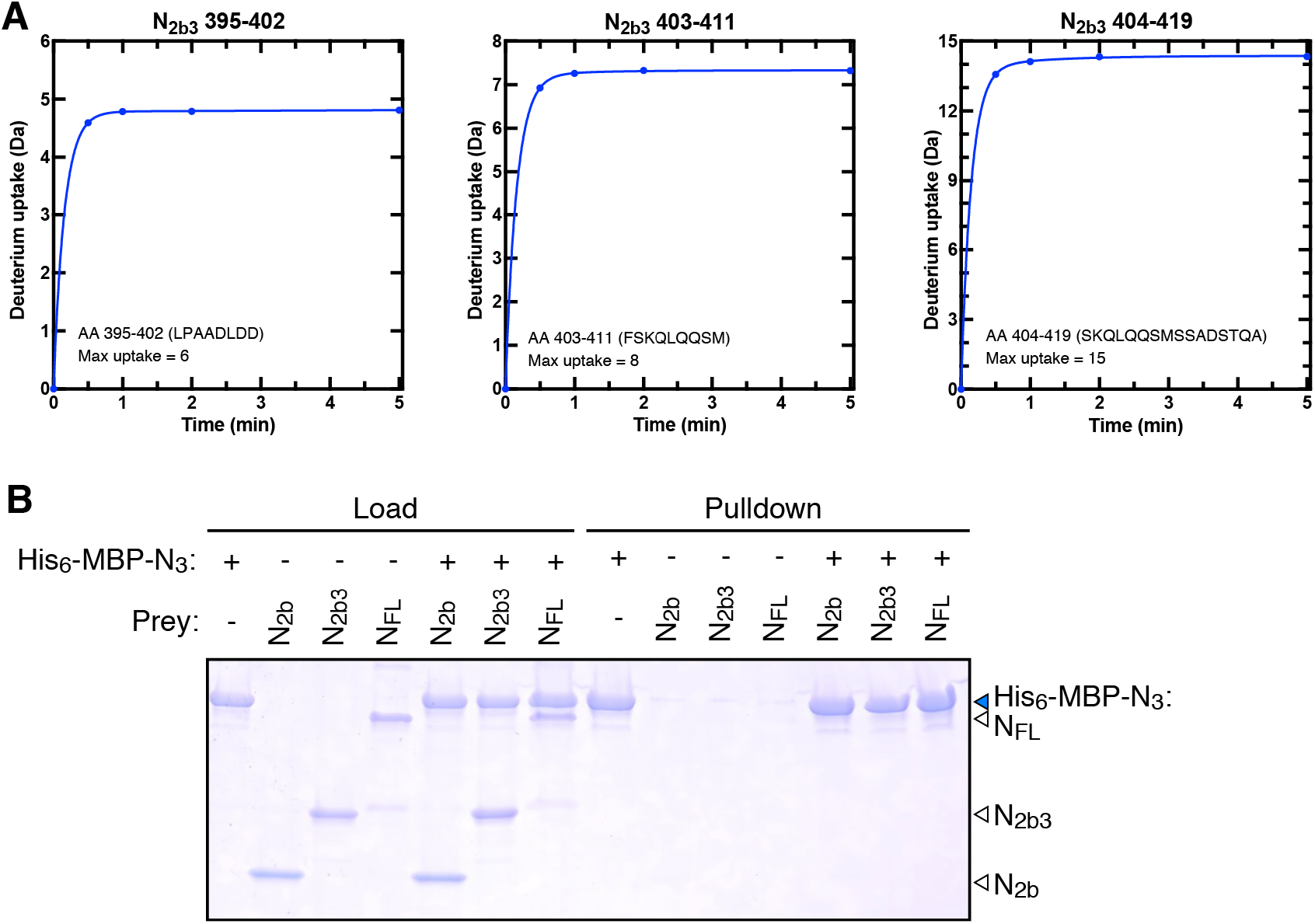
Role of the spacer B/N3 region in N protein self-assembly. (A) Absolute deuterium uptake plots for three peptides in the extreme C-terminus of N_2b3_, as shown in **Figure 4E**. (B) Nickel-NTA pulldown assay testing binding of His_6_-MBP-N_3_ with untagged N_2b_, N_2b3_, or N_FL_. Lanes 1-7: 10% of input (load); Lanes 8-14: Nickel-NTA bound (pulldown). His_6_-MBP-N_3_ does not strong bind any of the three prey proteins in this assay.

## Supplemental Tables

**Table S1.** Crystallographic data collection and refinement.

|  | N<sub>2b</sub> | His<sub>6</sub>-N<sub>2b</sub> |
| --- | --- | --- |
| **Data collection** |  |  |
| Synchrotron/Beamline | APS 24ID-E | APS 24ID-E |
| Date collected | May 12, 2020 | May 12, 2020 |
| Resolution (Å) | 66 – 1.42 | 74 – 1.45 |
| Wavelength (Å) | 0.97918 | 0.97918 |
| Space Group | P1 | P2<sub>1</sub> |
| Unit Cell Dimensions (a, b, c) Å | 43.72, 50.06, 69.34 | 71.15, 43.55, 74.23 |
| Unit cell Angles (α,β,γ) ° | 106.50, 90.09, 97.14 | 90, 94.87, 90 |
| I/σ (last shell) | 16.8 (3.0) | 16.0 (2.0) |
| <sup>1</sup>R<sub>sym</sub> (last shell) | 0.039 (0.310) | 0.058 (0.802) |
| <sup>2</sup>R<sub>meas</sub> (last shell) | 0.046 (0.353) | 0.063 (0.877) |
| <sup>3</sup>CC<sub>1/2</sub> (last shell) | 0.999 (0.957) | 0.998 (0.796) |
| Completeness (last shell) % | 91.1 (75.0) | 98.8 (91.9) |
| Number of reflections | 352232 | 528562 |
| *unique* | 96446 | 80163 |
| Multiplicity (last shell) | 3.7 (3.6) | 6.6 (6.0) |
| **Refinement** |  |  |
| Resolution (Å) | 66 – 1.42 | 74 – 1.45 |
| No. of reflections | 96393 | 80120 |
| *working* | 91660 | 76201 |
| *free* | 4733 | 3919 |
| <sup>4</sup>R<sub>work</sub> (last shell) (%) | 15.73 (25.33) | 16.73 (28.88) |
| <sup>4</sup>R<sub>free</sub> (last shell) (%) | 17.32 (24.32) | 18.02 (30.96) |
| **Structure/Stereochemistry** |  |  |
| No. of atoms | 7526 | 7768 |
| *hydrogen* | 3410 | 3580 |
| *solvent* | 618 | 493 |
| *ligand* | 0 | 30 |
| r.m.s.d. bond lengths (Å) | 0.007 | 0.009 |
| r.m.s.d. bond angles (°) | 0.888 | 0.922 |
| <sup>5</sup>SBGrid Data Bank ID | 785 | 786 |
| <sup>6</sup>Protein Data Bank ID | 6WZO | 6WZQ |

<sup>1</sup>R<sub>sym</sub> = $\sum_j |I_j - \langle I \rangle| / \sum_j I_j$ , where $I_j$ is the intensity measurement for reflection $j$ and $\langle I \rangle$ is the mean intensity for multiply recorded reflections.
<sup>2</sup>R<sub>meas</sub> = $\sum_h [v(n/(n-1)) \sum_j |I_{hj} - \langle I_h \rangle|] / \sum_h \langle I_h \rangle$
where $I_{hj}$ is a single intensity measurement for reflection $h$ , $\langle I_h \rangle$ is the average intensity measurement for multiply recorded reflections, and $n$ is the number of observations of reflection $h$ .
<sup>3</sup>CC<sub>1/2</sub> is the Pearson correlation coefficient between the average measured intensities of two randomly-assigned half-sets of the measurements of each unique reflection. CC<sub>1/2</sub> is considered significant above a value of ~0.15.
<sup>4</sup>R<sub>work, free</sub> = $\sum ||F_{obs}| - |F_{calc}|| / |F_{obs}|$ , where the working and free $R$ -factors are calculated using the working and free reflection sets, respectively.
<sup>5</sup>Diffraction data for each structure have been deposited with the SBGrid Data Bank (https://data.sbgrid.org) with the noted accession codes.
<sup>6</sup>Coordinates and structure factors for each structure have been deposited with the Protein Data Bank (http://www.pdb.org) with the noted accession codes.

**Table S2. N protein variants in SARS-CoV-2 patient sequences**

*(see attached Excel spreadsheet)*

